# PAK3 downregulation induces cognitive impairment following cranial irradiation

**DOI:** 10.1101/2023.06.30.547296

**Authors:** Haksoo Lee, Hyunkoo Kang, Changjong Moon, BuHyun Youn

## Abstract

Cranial irradiation is used for prophylactic brain radiotherapy as well as treatment of primary brain tumors. Despite its high efficiency, it often induces unexpected side effects, including cognitive dysfunction. Herein, we observed that mice exposed to cranial irradiation exhibited cognitive dysfunction, including altered spontaneous behavior, decreased spatial memory, and reduced novel object recognition. Analysis of actin cytoskeleton revealed that ionizing radiation (IR) disrupted the filamentous/globular actin (F/G-actin) ratio and downregulated the actin turnover signaling pathway p21-activated kinase 3 (PAK3)-LIM kinase 1 (LIMK1)-cofilin. Furthermore, we found that IR could upregulate microRNA-206-3p (miR-206-3p) targeting PAK3. As the inhibition of miR-206-3p through antagonist (antagomiR), IR-induced disruption of PAK3 signaling is restored. In addition, intranasal administration of antagomiR-206-3p recovered IR-induced cognitive impairment in mice. Our results suggest that cranial irradiation-induced cognitive impairment could be ameliorated by regulating PAK3 through antagomiR-206-3p, thereby affording a promising strategy for protecting cognitive function during cranial irradiation, and promoting quality of life in patients with radiation therapy.

## Introduction

Radiotherapy is an important component of cancer treatment, which utilizes high-energy radiation to damage the DNA of cancer cells and prevent their growth and division **(Brown et al., 2018; Owonikoko et al., 2014)**. Over the years, radiotherapy has undergone significant developments to improve its efficacy and minimize its side effects. Nevertheless, some adverse effects of radiotherapy have been reported, including cognitive impairment **(Houillier et al., 2019; Krull et al., 2018; Rapp et al., 2015; Shah et al., 2015)**. One type of radiotherapy that has been associated with such adverse effects is prophylactic cranial irradiation (PCI), which is used to prevent the spread of cancer to the brain and is commonly administered to patients with certain types of cancer, such as small-cell lung cancer and acute lymphoblastic leukemia, who are at high risk of developing brain metastasis **(Conklin et al., 2012; Xue et al., 2022)**. Notably, in the case of small-cell lung cancer, the cumulative risk of brain metastasis within 1 year was 14.6% in the PCI group and 40.4% in the non-PCI group. Accordingly, the 1-year survival rate was 27.1% in the PCI group and 13.3% in the non-PCI group (ClinicalTrials.gov number, NCT00016211, **(Slotman et al., 2007)**). However, it has been reported that the side effects of PCI inevitably affect 50–90% of long-term survivors with permanent and substantial cognitive impairment **(Greene-Schloesser et al., 2013; Makale et al., 2017; Rosi et al., 2008; Zhang et al., 2018)**. The survivors treated with whole-brain irradiation exhibit severe impairment in memory (54.9%), processing speed (66.3%), and executive function (68.3%) **(Brinkman et al., 2016; Krull et al., 2013)**. Moreover, the mechanisms underlying PCI-induced side effects remain poorly understood, and no treatment is currently available to address these adverse effects. Understanding the adverse effects associated with PCI is critical for developing strategies to minimize its impact on the cognitive function of cancer survivors.

Cognition is a complex mental process that involves the acquisition, processing, and application of knowledge **(Harvey, 2019; Jessen et al., 2020)**. The frontal cortex and hippocampus are two critical regions of the brain that play a key role in cognition, including memory, attention, and decision-making **(Christensen et al., 2022; Hogeveen et al., 2022; Price & Duman, 2020; Sakimoto et al., 2021)**. Cognitive impairment is a common feature of many psychiatric diseases, such as schizophrenia **(Gebreegziabhere et al., 2022; Jauhar et al., 2022)** and major depressive disorder **(Douglas et al., 2018; Wen et al., 2022)**, and can significantly impact an individual’s quality of life. Despite the development of numerous therapies for psychiatric diseases, the limitations of present cure have resulted in cognitive impairment remaining a persistent and unmet need in treating these conditions. Therefore, a deeper understanding of the underlying mechanisms of cognitive impairment is needed to develop more effective treatments and improve patients’ quality of life suffering from these disorders.

Cognition is closely associated with dendritic spine morphology, which plays a key role in synaptic plasticity in the frontal cortex and hippocampus **(Delevich et al., 2020; Lamprecht, 2021; Yang et al., 2021)**. Dendritic spines are small protrusions that extend from the dendrites of neurons and play a critical role in synaptic plasticity, which is the ability of synapses (the junctions between neurons) to change strength and structure in response to experience **(Berry & Nedivi, 2017)**. Synaptic plasticity is believed to underlie learning and memory processes. The morphology of dendritic spines, which includes their shape, size, and density, is closely linked to synaptic plasticity and can influence cognitive function.**(Huang et al., 2021; Nakahata & Yasuda, 2018)** For example, a higher density of dendritic spines in the frontal cortex has been associated with better cognitive function **(Counts et al., 2006; Delevich et al., 2020)**, while a decrease in dendritic spine density in the hippocampus has been linked to memory impairment **(Bosch et al., 2014; Kandimalla et al., 2018; Szatmari et al., 2021)**. In neurons, the dendritic spine morphology is closely related to the balance between two forms of the cytoskeletal protein actin: filamentous actin (F-actin) and globular actin (G-actin) **(Hlushchenko et al., 2016; Lei et al., 2016)**. F-actin is a rigid, filament-like structure that provides the structural stability required for dendritic spine maturation and synaptic function. G-actin is a monomeric form of actin that serves as the building block for F-actin and is critical for actin dynamics and the formation of new dendritic spines. An appropriate balance between F-actin and G-actin is essential for maintaining dendritic spine structure and function **(Hlushchenko et al., 2016; Lei et al., 2016)**. Disruptions in this balance have been linked to cognitive dysfunction in various neurological and psychiatric disorders.

p21-activated kinase 3 (PAK3) is a serine/threonine kinase that plays a critical role in the regulation of dendritic spine morphology and synaptic plasticity **(Combeau et al., 2012; Thévenot et al., 2011)**. PAK proteins function as Rac1 and Cdc42-specific effector molecules that mediate cytoskeletal remodeling by regulating LIM kinase (LIMK), oncoprotein 18 (OP18), and tubulin folding cofactor b (TBCB) **(Molli et al., 2009)**. PAK3 is highly expressed in the brain, particularly in postmitotic neurons of the cerebral cortex and the hippocampus, where it shows a diffuse distribution throughout the soma and proximal dendrites. PAK3-deficient mice exhibit defects in the morphology of dendritic spines, which lead to impairments in synaptic plasticity and cognitive function **(Meng et al., 2005)**. Several studies have reported that PAK3 is implicated in cognitive impairment in various neurological and psychiatric disorders, including schizophrenia and Alzheimer’s disease (AD) **(Lauterborn et al., 2020; Morrow et al., 2008)**. However, the mechanism underlying PAK3-associated cognitive impairment has not yet been fully understood.

In this study, we investigated the impact of cranial irradiation on cognitive function and its underlying mechanisms. Our results demonstrate that ionizing radiation (IR) reduced the level of PAK3, leading to downregulation of PAK3-LIMK1-cofilin signaling, which correlates with the maturation of dendritic spines. We identified that microRNA-206-3p (miR-206-3p) regulates PAK3 and that its antagonist, antagomiR-206-3p, restores PAK3 levels and cognitive function in irradiated mouse models. Our study provides valuable insights into the impact of cranial irradiation on the brain and its potential mechanisms, and a promising strategy to prevent PCI-induced cognitive dysfunction.

## Results

### Cranial irradiation induces cognitive impairment in mice

Given that cranial irradiation is known to induce cognitive impairment **(Ding et al., 2022; Krull et al., 2018; Makale et al., 2017)**, we first evaluated whether cranial irradiation impaired cognitive function in mice. To assess the effect of IR on cognitive function, we exposed mice to IR. Subsequently, we performed behavioral tests following the overall schedule (Fig. 1A). A total dose of 60 Gy in 30 fractions is typically delivered to the brain in patients with PCI **(Suwinski, 2021; Wegner et al., 2019)**, and previous studies reported that a total dose of 10 Gy caused impaired neurogenesis and cognitive behavior in mice **(Pazzaglia et al., 2020; Raber et al., 2004)**. In this regard, mice were irradiated with five fractions of 2 Gy administered to the cranial region **(Ceyzériat et al., 2021; Wilson et al., 2020)**. Individual behavioral tests, such as the open field test (OPF), Y-maze, and novel object recognition test (NOR), were performed from days 3 to 8 post-IR. In mice treated with IR, spontaneous alteration and time in the novel arm were significantly reduced by 66.67 and 58.14%, respectively (Fig. 1B and 1C). The time spent with objects did not differ between familiar and novel objects in IR-treated mice (Fig. 1C). Our previous study has shown that IR can induce depression-like behavior **(Kim et al., 2022)**. IR-treated mice exhibited reduced time in the center of the OPF, thereby indicating depressive-like behavior (Fig. S1A). Based on the findings in the OPF, IR partially induced depression-like behavior, but the total distance traveled did not differ. These results indicate that IR could impair cognitive function and that depressive-like behavior does not impact overall moving behavior.

**Fig 1.**
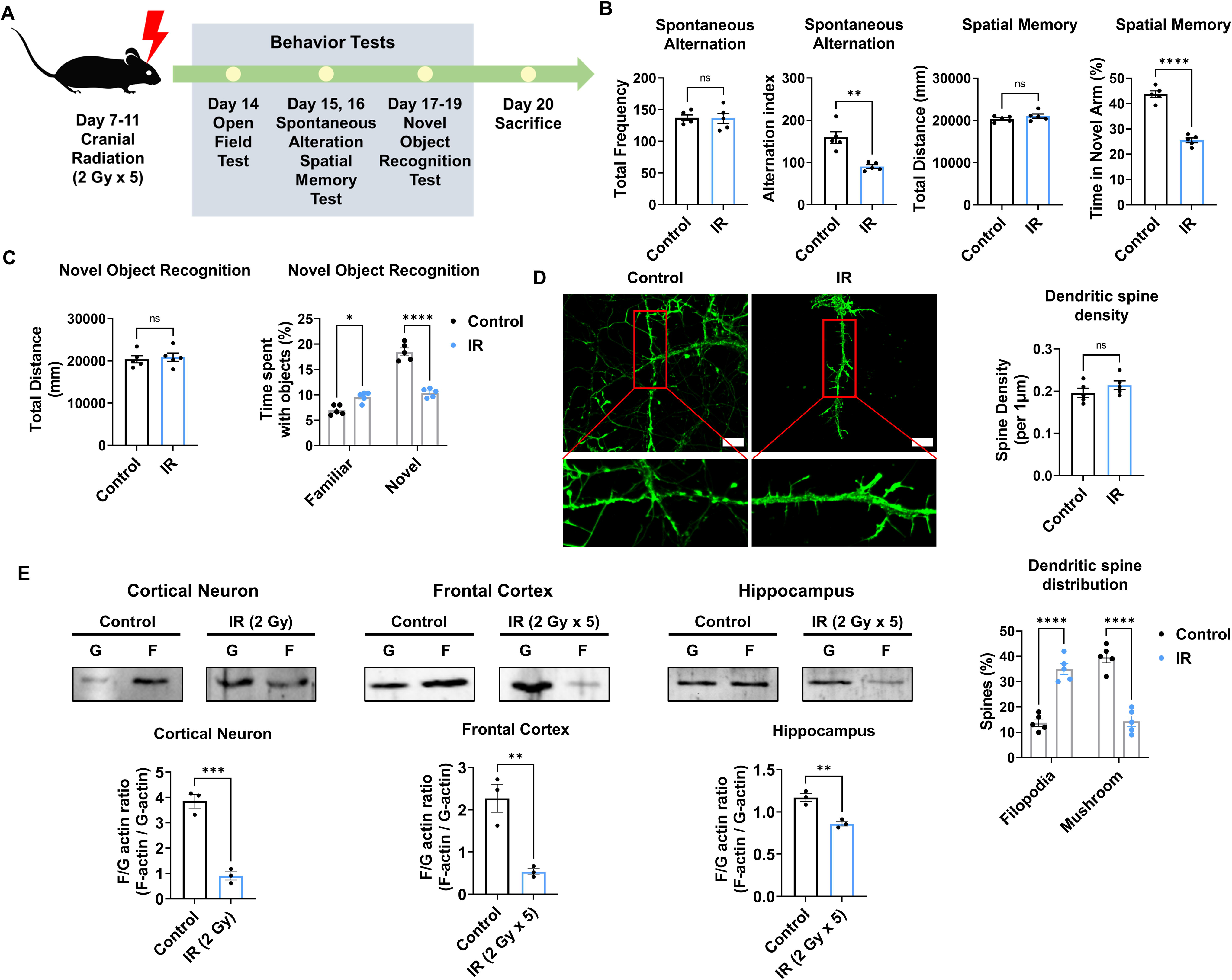
Cranial irradiation affects cognitive function and the maturation of dendritic spine. **(A)** A scheme illustrating the schedule of ionizing radiation (IR) and overall behavioral tests. (**B)** The results of spontaneous alteration (left) and spatial memory test (right) after IR using Y-maze. The parameters of spontaneous alteration and spatial memory are described in methods. (**C)** The result of novel object recognition test after IR. (**D)** Left: representative fluorescence image of filamentous actin (F-actin, green) after IR in cortical neuron. Right: The density and distribution of dendritic spine in IR group compared to control group. Scale bars, 10 μm. **(E)** The alterations of F/G-actin ratio after IR in cortical neuron, frontal cortex, and hippocampus. Statistical analysis was performed with Student’s t test for (**B**, left of **C**, and **E**) and one-way ANOVA plus a Tukey’s multiple comparisons test for (right of **C** and **D**). ns, non-significant; *p < 0.05; **p < 0.01; ***p < 0.001; ****p < 0.0001.

Synapse formation on dendrites is known to impact cognitive behavior **(Arendt, 2009; Baloyannis, 2009)**. Therefore, we examined changes in the dendritic spines of the cortical neurons after IR exposure. Cortical neurons were treated with a single 2 Gy at 14 days *in vitro* (DIV). Dendritic spine density did not differ between the non-IR and IR groups. In IR-treated neurons, mature dendritic spines (mushroom) were significantly decreased by 37.5%, whereas immature dendritic spines (filopodia) were increased by 37.14% (Fig. 1D). The maturation of dendritic spines depends on the F/G-actin ratio **(Chen et al., 2007; Rex et al., 2009)**. Accordingly, we evaluated whether IR can impact the F/G-actin ratio in neurons. Exposure to IR decreased the F/G-actin ratio in cortical neurons (21.05%), frontal cortex (21.74%), and hippocampus (73.91%) (Fig. 1E). Overall, these results indicated that IR could induce cognitive impairment by disrupting dendritic spine maturation.

### IR decreases PAK3-LIMK1-cofilin signaling

To identify the molecular mechanism of IR-induced dendritic spine alteration, we analyzed microarray data from the brains of mice treated or un-treated with cranial radiation, which was generated in a previous study (GSE94440) **(Kang et al., 2018)**. Among the 15 coding genes that were up- and downregulated by cranial irradiation, we focused on the PAK3 gene, which showed a 0.5-fold altered expression in the irradiated brain, considering both the “synapse organization” and the “dendritic spine development” ontologies (Fig. 2A). In addition, PAK3 is mainly expressed in the frontal cortex and hippocampus, both crucial regions for cognitive function in the brain (Fig. S2A) **(Christensen et al., 2022; Price & Duman, 2020)**. As shown in Fig. 2b, IR decreased mRNA and protein levels of PAK3 in cortical neurons (49.48% and 44.30%, respectively). In addition, IR decreased mRNA level of PAK3 in frontal cortex (61.67%) and hippocampus (66.67%) (Fig. S2B). Based on immunocytochemistry data, PAK3 was localized on dendrites in cortical neurons (Fig. 2C). These results suggested that PAK3, which is correlated with synapses and dendritic spines, is a candidate for IR-induced dendritic spine alteration.

**Fig 2.**
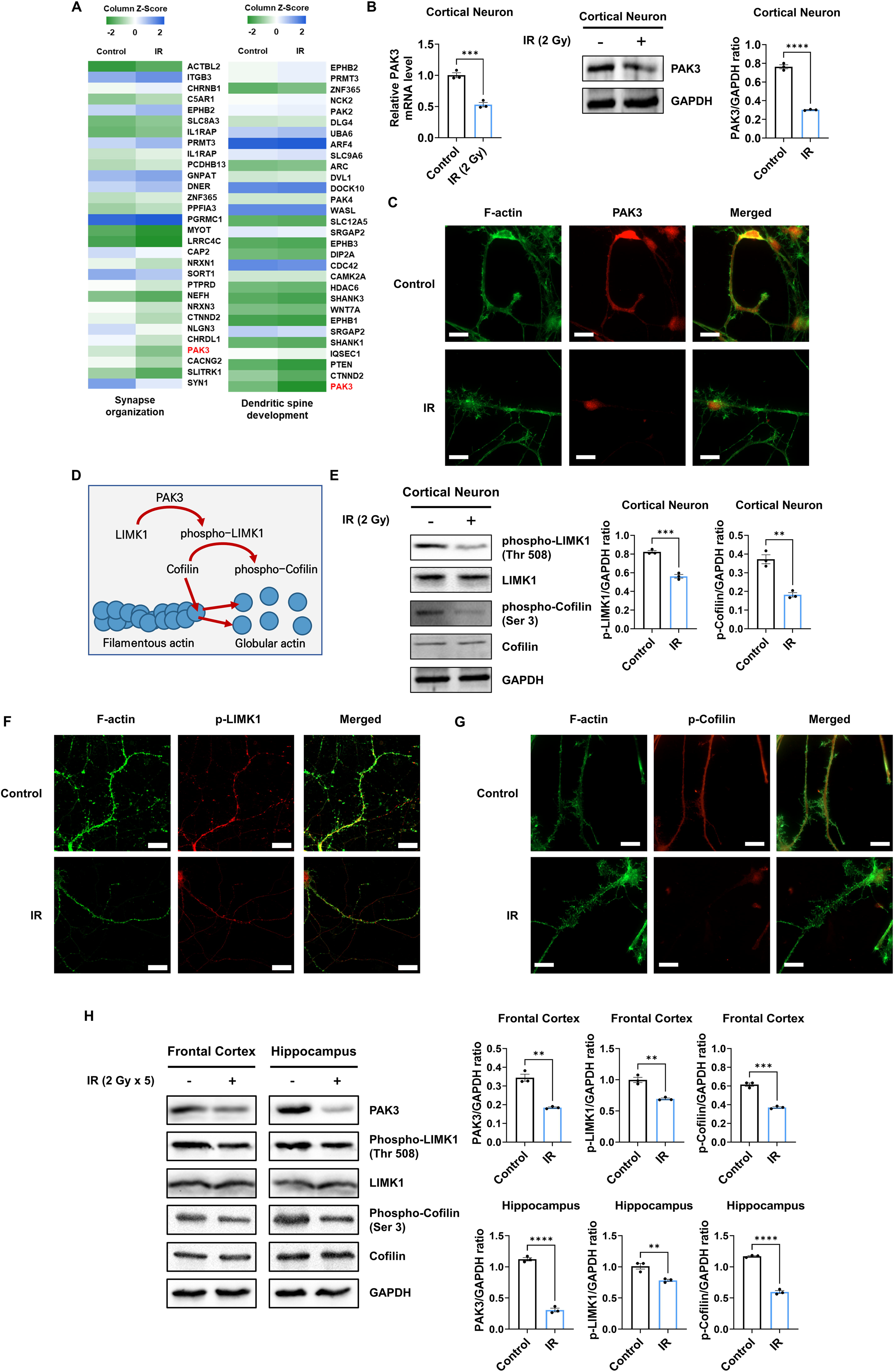
IR decreases phosphorylation of LIMK1 and Cofilin via PAK3 downregulation. **(A)** The heatmap for the expression of genes in the term ‘‘synapse organization’’ and “dendritic spine development” in the irradiated brain. (**B)** The changes of PAK3 mRNA and protein levels after IR in cortical neuron. Each western blot bands are quantified by ImageJ. (**C)** Representative fluorescence image of F-actin (green) and PAK3 (red) in cortical neuron. Scale bars, 10 μm. (**D)** Schematic illustration of PAK3-LIMK1-cofilin signaling. (**E)** Left: the protein levels of phosphorylated LIMK1, LIMK1, phosphorylated cofilin, and cofilin after IR in cortical neuron. Right: each western blot bands are quantified by ImageJ. (**F**-**G)** Representative fluorescence image of F-actin (green) and phosphorylated LIMK1 (**F**, red) or phosphorylated cofilin (**G**, red) in cortical neuron. Scale bars, 10 μm. (**H**) Left: the protein levels of phosphorylated LIMK1, LIMK1, phosphorylated cofilin, and cofilin after IR in frontal cortex and hippocampus. Right: each western blot bands are quantified by ImageJ. Statistical analysis was performed with Student’s t test. ns, non-significant; **p < 0.01; ***p < 0.001; ****p < 0.0001.

Given that PAK3 inhibits severing and promotes the branching of F-actin **(Rane & Minden, 2014)**, we further examined actin dynamics-related PAK3 signaling. PAK3 phosphorylates LIMK1 to activate it, and activated LIMK1, in turn, phosphorylates cofilin to its inactive state. Activated cofilin promotes actin turnover, while inactivated cofilin suppresses actin turnover (Fig. 2D) **(Bamburg et al., 2021; Edwards et al., 1999)**. Therefore, we examined whether IR can affect the activation of LIMK1 and cofilin. Herein, we observed that IR decreased the expression of phosphorylated LIMK1 and cofilin in cortical neurons, frontal cortex, and hippocampus (Fig. 2E and H). In addition, phosphorylated LIMK1 and cofilin were less expressed and localized on the dendrites in cortical neurons (Fig. 2F and G). These results suggested that IR could decrease the activation and localization of LIMK1 and cofilin in dendritic spines by downregulating PAK3 expression. However, IR did not alter other downstream factors related to cytoskeleton regulation, such as oncoprotein 18 (OP18) and tubulin-folding cofactor B (TBCB) (Fig. S3A and S3B). Overall, these results showed that IR decreased the expression and localization of PAK3, which, in turn, decreased the phosphorylation of LIMK1 and cofilin.

### PAK3 regulates dendritic spine morphology and maturation

Given that PAK3 is associated with synapse organization and dendritic spine development, we further examined whether the regulation of PAK3 could impact dendritic spines. First, we knocked down PAK3 using a short hairpin RNA (shRNA). Every two shRNAs targeting PAK3 successfully reduced the mRNA level of PAK3 by approximately 30% (Fig. 3A). We employed shRNA #1 to examine the effects of PAK3 downregulation. Downregulated PAK3 expression reduced the F/G-actin ratio (Fig. 3B) and LIMK1-cofilin signaling (Fig. 3C). Dendritic spines were observed after PAK3 downregulation. We noted that the dendritic spine density did not differ between the control and PAK3 downregulation groups. However, the number of mushroom spines was lower than that of filopodia following PAK3 downregulation (Fig. 3D). Compared with the filopodia spine, the mushroom spine is a more mature form that typically transmits nerve signals to other neurons **(Hering & Sheng, 2001)**. We examined the effects of PAK3 overexpression (Fig. 3E). PAK3 overexpression affects the expression and phosphorylation of LIMK1 and cofilin. We evaluated the recovery of IR-induced downregulated PAK3 signaling after PAK3 overexpression. Notably, PAK3 overexpression could restore IR-induced downregulation of F/G-actin (Fig. 3F). Following alterations in the F/G-actin ratio, mushroom spines were more abundant than filopodia in IR-treated PAK3 overexpressing neurons (Fig. 3G). These results suggested that PAK3 could regulate the F/G-actin ratio and maturation of dendritic spine through LIMK1-cofilin signaling. Considering IR-induced decreased PAK3 signaling, upregulation of PAK3 expression could restore the F/G-actin ratio and dendritic spine distribution to normal levels in murine neurons.

**Fig 3.**
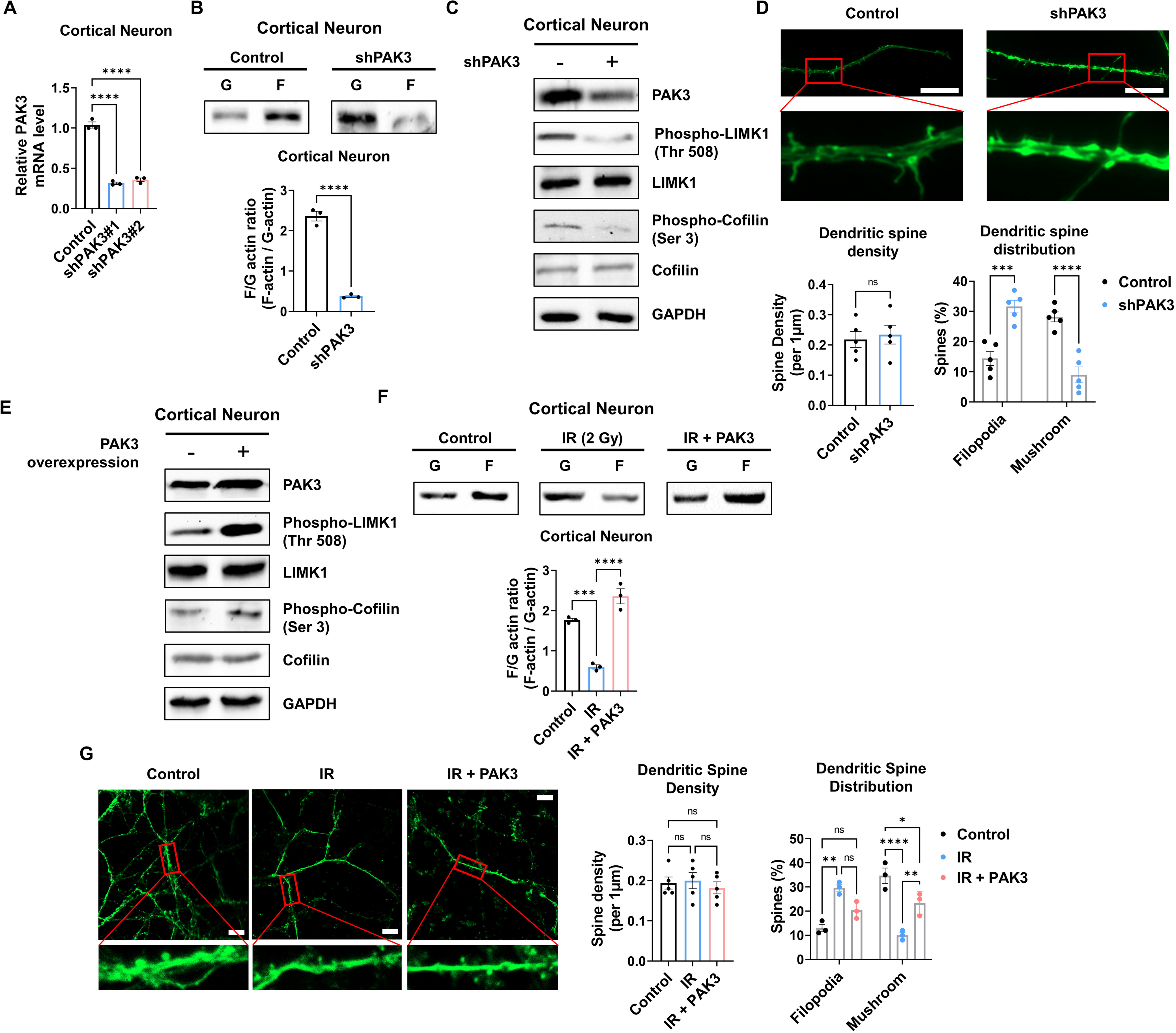
PAK3 regulates F/G-actin and dendritic spine in neuron. **(A)** The changes of PAK3 mRNA levels after PAK3 downregulation (shPAK3) in cortical neuron. (**B)** The F/G-actin ratio after expression of PAK3 downregulation (shPAK3) in cortical neuron. Each F-actin and G-actin are quantified by ImageJ. (**C)** The protein levels of phosphorylated LIMK1, LIMK1, phosphorylated cofilin, and cofilin after PAK3 downregulation (shPAK3) in cortical neuron. (**D)** Upper: representative fluorescence image of filamentous actin (F-actin, green) after PAK3 downregulation (shPAK3) in cortical neuron. Lower: the density and distribution of dendritic spine in shPAK3 group compared to control group. Scale bars, 10 μm. **(E)** The protein levels of PAK3, phosphorylated LIMK1, LIMK1, phosphorylated cofilin, and cofilin after PAK3 overexpression in cortical neuron. (**F)** The F/G-actin ratio after IR or IR with PAK3 overexpression in cortical neuron. Each F-actin and G-actin are quantified by ImageJ. (**G)** Upper: representative fluorescence image of filamentous actin (F-actin, green) after IR and/or PAK3 overexpression in cortical neuron. Lower: the density and distribution of dendritic spine in control, IR and/or PAK3 overexpression group. Scale bars, 10 μm. Statistical analysis was performed with Student’s t test for (**B** and **D**) and one-way ANOVA plus a Tukey’s multiple comparisons test for (**A**, **D**, **F**, and **G**). ns, non-significant; *p < 0.05; **p < 0.01; ***p < 0.001; ****p < 0.0001.

To explore whether IR can decrease PAK3 signaling and dendritic spine maturation in human neurons, we differentiated neural progenitor cells into neurons. Herein, we observed that the expression of stemness markers (SOX2, Nestin, and PAX6) was decreased in differentiated neurons (Fig. S4A). In addition, the expression of neuronal (NeuN) and synaptic (PSD95) markers was increased in differentiated neurons when compared with that in neural progenitor cells (Fig. S4B). The altered expression of markers suggests that human neural progenitor cells can differentiate into human neurons. Therefore, differentiated neurons were used to confirm alterations in human neurons. IR decreased neuronal viability in human differentiated neurons, with approximately 80% survival (Fig 4A). However, IR did not alter the mature neuronal marker, NeuN (Fig S5A). These results indicate that IR-induced disruption of PAK3 signaling occurs in surviving neurons following irradiation. Consistent with previous murine neuron data, IR reduced the F/G-actin ratio (Fig. 4B). Based on IR-induced altered F/G-actin ratio, we hypothesized that IR could reduce PAK3-LIMK1-cofilin signaling in differentiated neurons. IR decreased the expression of PAK3, phosphorylated-LIMK1, and phosphorylated-cofilin in human neurons (Fig. 4C and D). A previous study has revealed that IR can reduce mature dendritic spines and enhance immature dendritic spines (Fig. 1D); IR downregulated the mature dendritic spine and upregulated the immature dendritic spine in human neurons (Fig. 4E). To confirm that these changes were caused by IR-induced PAK3 downregulation, we used shRNA to downregulate PAK3 expression in differentiated neurons. Consistent with the IR effect on dendritic spines, downregulated PAK3 expression increased the number of immature dendritic spines (Fig. 4F). Collectively, PAK3 signaling is essential for the maturation of dendritic spines, and IR could disrupt PAK3 signaling in human neurons.

**Fig 4.**
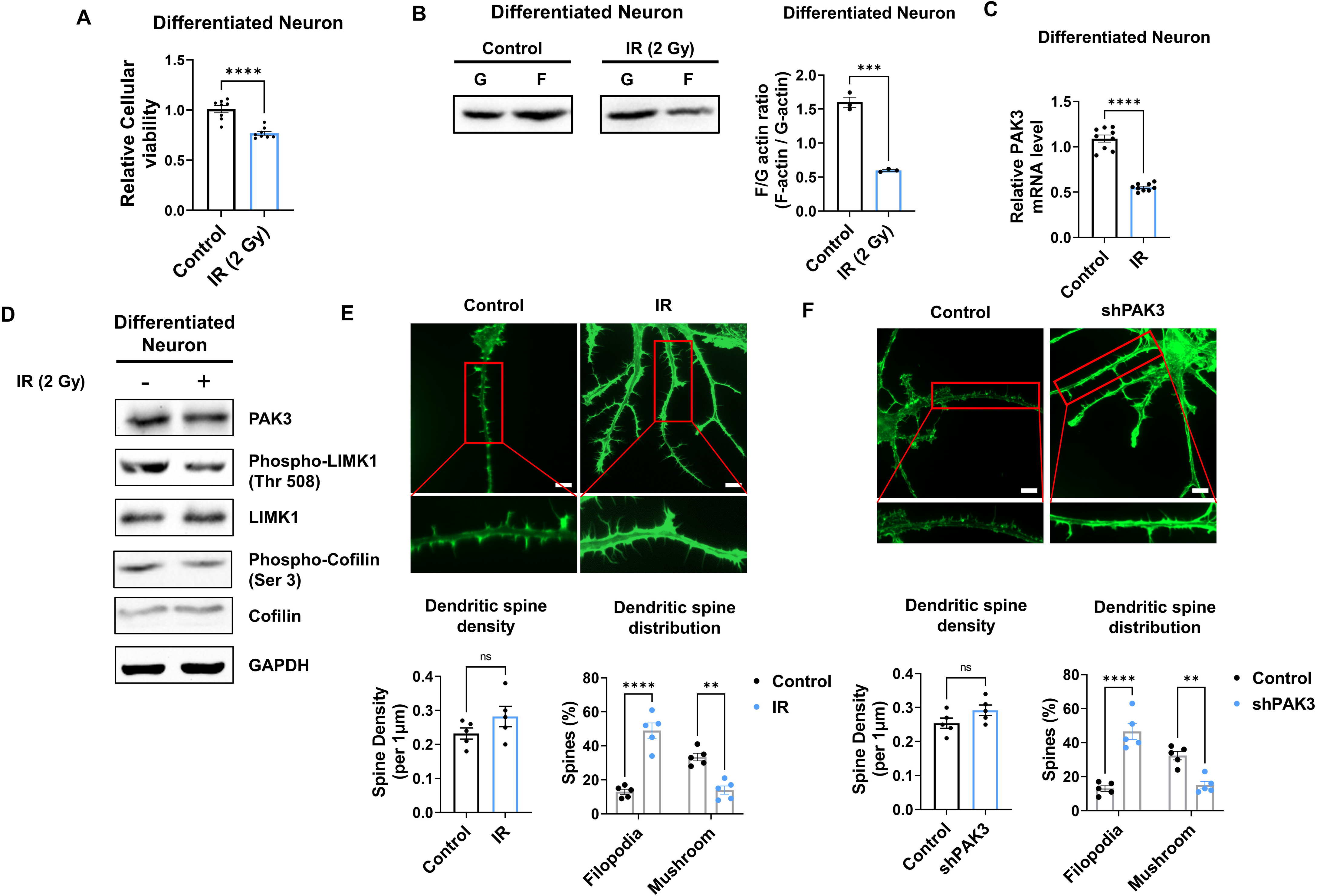
IR decreases PAK3-LIMK1-Cofilin signaling in differentiated human neuron. **(A)** The F/G-actin ratio after IR in differentiated neuron. Each F-actin and G-actin are quantified by ImageJ. (**B)** The mRNA level of PAK3 after IR in differentiated neuron. (**C)** The protein levels of PAK3, phosphorylated LIMK1, LIMK1, phosphorylated cofilin, and cofilin after IR in differentiated neuron. (**D)** Upper: representative fluorescence image of filamentous actin (F-actin, green) after IR in differentiated neuron. Lower: the density and distribution of dendritic spine in IR group compared to control group. Scale bars, 10 μm. (**E)** Upper: representative fluorescence image of filamentous actin (F-actin, green) after PAK3 downregulation (shPAK3) in differentiated neuron. Lower: the density and distribution of dendritic spine in shPAK3 group compared to control group. Scale bars, 10 μm. Statistical analysis was performed with Student’s t test for (**A**, **B**, **D**, and **E**) and one-way ANOVA plus a Tukey’s multiple comparisons test for (**D** and **E**). ns, non-significant; **p < 0.01; ***p < 0.001; ****p < 0.0001.

### IR-induced miR-206-3p decreases PAK3 expression, and its inhibition rescued PAK3 signaling

To elucidate the mechanism underlying IR-induced changes in PAK3 expression, we performed a luciferase reporter assay for the PAK3 promoter. In murine neurons, IR failed to induce any change in luciferase activity (Fig. 5A), suggesting that IR does not induce PAK3 expression by regulating transcription levels. Next, we hypothesized that IR might alter the miRNAs targeting PAK3. Using miRNA databases (miRDB, TargetScan, and TarBase), we analyzed miRNAs targeting PAK3, selecting three miRNA candidates (Fig. 5B). Among the three miRNAs, IR significantly upregulated only miR-206-3p in both mice and humans (Fig. 5C). To validate whether IR can impact miR-206-3p expression and examine the effects of miR-206-3p inhibition, we determined miR-206-3p levels after IR and treatment with an inhibitor of miR-206-3p (antagomiR-206-3p). IR increased miR-206-3p expression in both mouse and human neurons (Fig. 5D). In addition, antagomiR-206-3p successfully inhibited IR-induced miR-206-3p expression in mouse and human neurons (Fig. 5D). Consistent with the miRNA inhibition data, IR-induced PAK3 downregulation was restored to approximately 90% of the control in both neurons (Fig. 5E). To confirm whether miR-206-3p could affect PAK3 signaling, we examined whether treatment with miR-206-3p could alter PAK3-LIMK1-cofilin signaling. As expected, miR-206-3p decreased PAK signaling, and the respective antagonist partially restored the downregulated PAK3 signaling (Fig. 5F and 5G). Additionally, the post-synaptic marker, PSD-95, was decreased by miR-206-3p treatment. However, a mature neuronal marker (NeuN) and non-neuronal markers (GFAP and IBA-1) were not alterd upon miR-206-3p treatment (Fig. S5A and S5B). In addition, consistent with the effects of antagomiR, IR disrupted the distribution of dendritic spines, and IR with antagomiR treatment restored dendritic spine maturation (Fig. 5H). These results suggested that IR-induced miR-206-3p decreased PAK3 signaling and dendritic spine maturation in mouse and human neurons.

**Fig 5.**
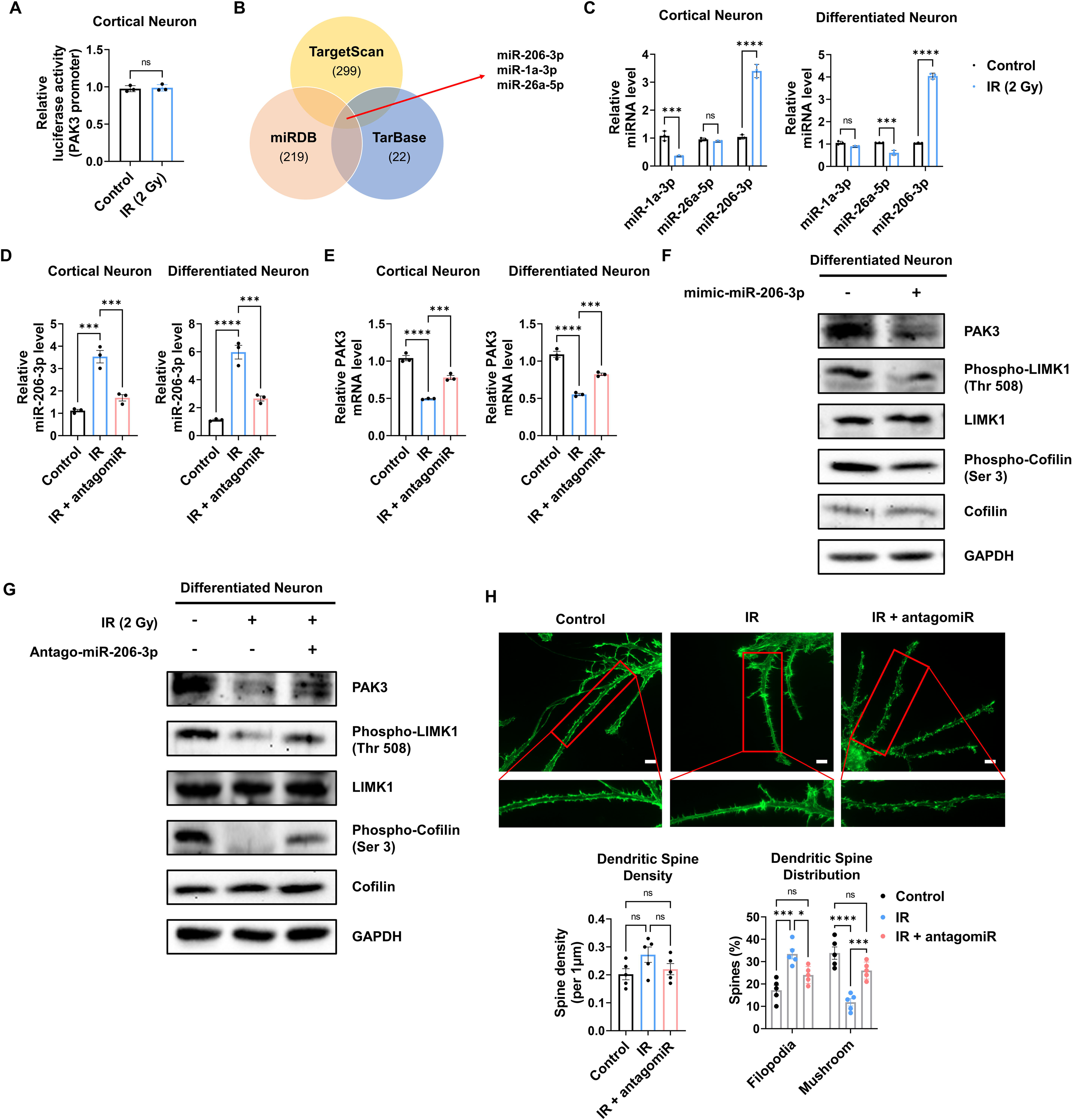
IR-induced miR-206-3p affects PAK3-LIMK1-cofilin signaling. **(A)** The relative luciferase activity of PAK3 promoter after IR in cortical neuron. (**B)** Venn diagram of predictable PAK3-targeting miRNAs from databases (miRDB, TargetScan, and TarBase). (**C)** The relative miRNA expression of 3 overlapped miRNAs after IR in cortical neuron and differentiated neuron. (**D)** The relative miRNA expression of miR-206-3p after IR and/or antagomiR-206-3p treatment in cortical neuron and differentiated neuron. (**E)** The mRNA level of PAK3 after IR and/or treatment of antagomiR-206-3p in cortical neuron and differentiated neuron. (**F)** The protein levels of PAK3, phosphorylated LIMK1, LIMK1, phosphorylated cofilin, and cofilin after treatment of miR-206-3p mimic in differentiated neuron. (**G)** The protein levels of PAK3, phosphorylated LIMK1, LIMK1, phosphorylated cofilin, and cofilin after IR and/or treatment of antagomiR-206-3p in differentiated neuron. (**H)** Upper: representative fluorescence image of filamentous actin (F-actin, green) after IR and/or treatment of antagomiR-206-3p in cortical neuron. Lower: the density and distribution of dendritic spine in control, IR and/or treatment of antagomiR-206-3p group. Scale bars, 10 μm. Statistical analysis was performed with Student’s t test for **A** and one-way ANOVA plus a Tukey’s multiple comparisons test for the others. ns, non-significant; ***p < 0.001; ****p < 0.0001.

### Intranasal administration of antagomiR-206-3p rescues cognitive function

Given that antagomiR for miR-206-3p effectively attenuated IR-mediated alterations in dendritic spines, we next inhibited miR-206-3p to recover cognitive impairment *in vivo*. To efficiently deliver antagomiR to the brain, antagomiR was administered intranasally. Intranasal administration is a less invasive method for drug delivery, easily accessible to adult and pediatric patients. According to the previous *in vivo* experimental protocol (Fig. 1A), mice were exposed to IR with five fractions (over a 24 h interval) of 2 Gy to the cranial region and subsequently treated with antagomiR via intranasal administration. Then, OPF, Y-maze, and NOR tests were performed from days 2–7 post-intranasal administration. Behavioral tests were performed according to the overall schedule (Fig. 6A). We then administered Cy5-labeled antagomiR to IR-treated mice via intranasal administration (Fig. 6B). To validate the delivery efficiency of antagomiR, we examined Cy5 fluorescence in the brain after intranasal administration (Fig. 6C). Treatment with Cy5-antagomiR treatment dose-dependently increased Cy5 fluorescence. As the frontal cortex and hippocampus are cognition-related central regions in the brain **(Christensen et al., 2022; Hogeveen et al., 2022; Price & Duman, 2020; Sakimoto et al., 2021)**, we confirmed IR and antagomiR effects in the frontal cortex and hippocampus. Consistent with *in vitro* results (Fig. 5D), antagomiR decreased the IR-induced miR-206-3p expression in the frontal cortex and hippocampus (Fig. 6D). IR-treated mice administered antagomiR exhibited attenuated spontaneous alteration, as well as time in the novel arm and time spent with the novel object (Fig. 6E and 6F). In addition, time spent in the center of the OPF was restored in IR-treated mice administered antagomiR (Fig. S6A and S6B). These results showed that antagomiR could successfully attenuate IR-induced cognitive impairment. To validate whether antagomiR-induced recovery of cognitive function is mediated via the restoration of PAK3 signaling to normal levels, we confirmed that antagomiR restored IR-induced downregulation of PAK3 signaling. Consistently, antagomiR treatment increased the expression of PAK3, phosphorylated-LIMK1, and phosphorylated-cofilin in the frontal cortex and hippocampus (Fig. 6G). In addition, the F/G-actin ratio in the frontal cortex and hippocampus was restored to normal when compared with that in the IR-treated group (Fig. 6H). Collectively, these results suggested that antagomiR could be an effective strategy for IR-induced cognitive impairment by restoring PAK3 signaling.

**Fig 6.**
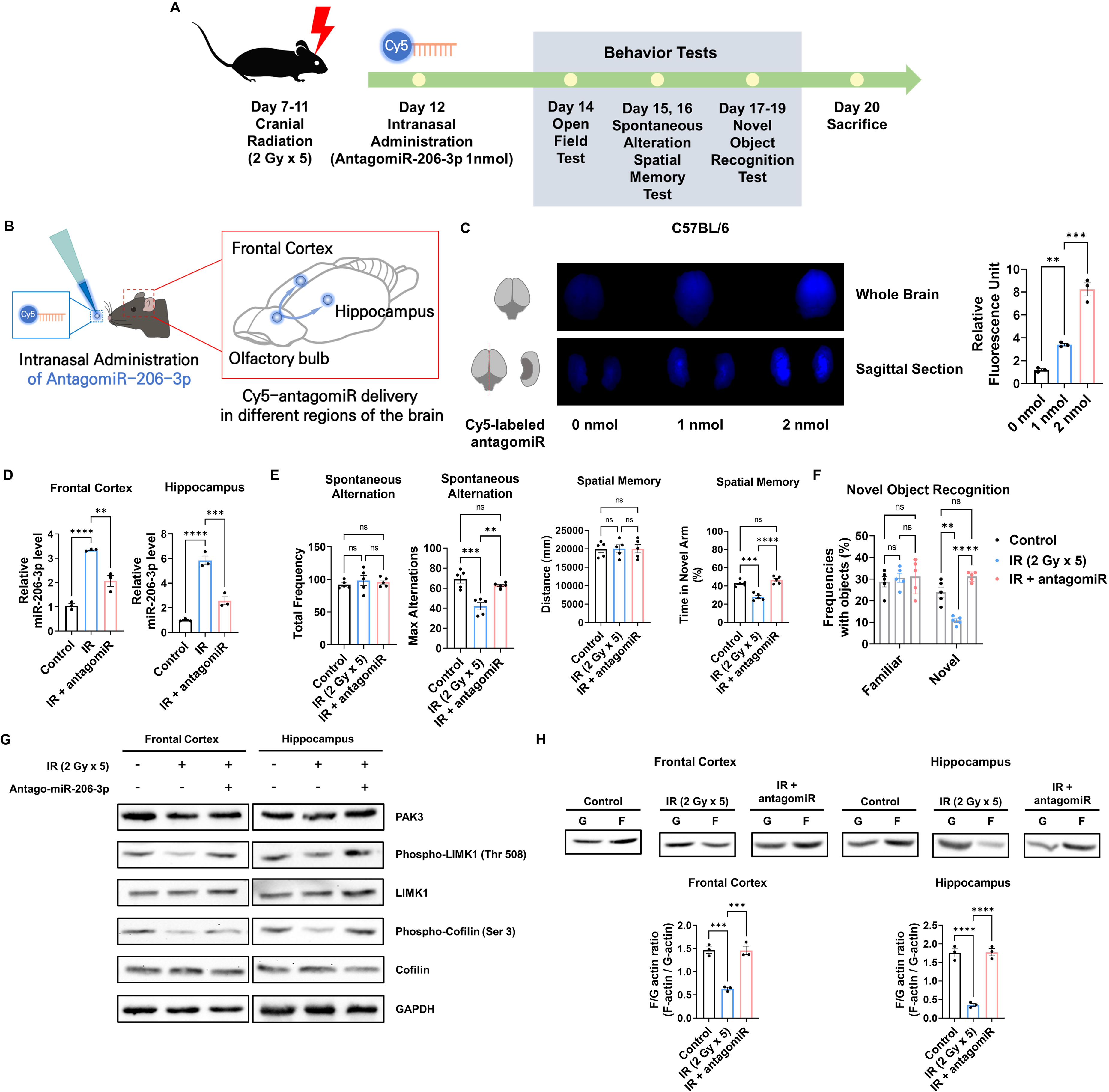
Intranasal administration of antagomiR-206-3p recovers PAK3 signaling and cognitive impairment after IR. **(A)** A scheme illustrating the schedule of IR and antagomiR-206-3p treatment and overall behavioral tests. (**B)** Schematic illustration of antagomir-206-3p delivery via intranasal administration. (**C)** The fluorescence images and relative units of cy5-antagomiR-206-3p in the brain of mice 24 h after intranasal administration. (**D)** The relative miRNA expression of miR-206-3p after IR and/or antagomiR-206-3p treatment in frontal cortex and hippocampus. (**E)** The results of spontaneous alteration (left) and spatial memory test (right) after IR and/or antagomiR-206-3p using Y-maze. The parameters of spontaneous alteration are described in methods. (**F)** The result of novel object recognition test after IR and/or antagomiR-206-3p. (**G)** The protein levels of PAK3, phosphorylated LIMK1, LIMK1, phosphorylated cofilin, and cofilin after IR and/or treatment of antagomiR-206-3p in frontal cortex and hippocampus. (**H)** The alterations of F/G-actin ratio after IR and/or treatment of antagomiR-206-3p in frontal cortex and hippocampus. Statistical analysis was performed with one-way ANOVA plus a Tukey’s multiple comparisons test. ns, non-significant; **p < 0.01; ***p < 0.001; ****p < 0.0001.

## Discussion

Radiotherapy is a common treatment for cancer to stop or prevent cancer cells from growing or spreading in the body. In particular, PCI is widely used for preventing brain metastasis, but it may cause side effects including cognitive impairment, which significantly impacts quality of life of patients. In this study, we investigated the effect of cranial irradiation on cognitive function and the underlying neuronal mechanisms using in vivo and in vitro models. Our results revealed that cranial irradiation reduced cognitive function and downregulated the expression of PAK3, a gene involved in dendritic spine development and synaptic plasticity in the frontal cortex and hippocampus. We demonstrated that the PAK3-LIMK1-cofilin signaling pathway was affected by IR, leading to dendritic spine maturation impairment. Additionally, we found that miR-206-3p regulated PAK3 expression and that antagomiR-206-3p treatment restored PAK3 levels and cognitive function in the irradiated mice. Our findings suggest that PAK3 may play a critical role in cognitive function after cranial irradiation, highlighting a potential therapeutic target to mitigate the adverse effects of radiotherapy (Fig. S7).

PAK3 expression is altered in several neurodegenerative diseases, suggesting that the decrease in PAK3 may be partly involved in cognitive or developmental symptoms. PAK3 mRNA levels are significantly reduced in the hippocampus of subjects affected with depression **(Fuchsova et al., 2016)**. Additionally, post-mortem brains of Down syndrome and AD patients, PAK3 proteins are markedly reduced with abnormal cofilin activation in neurites **(Lauterborn et al., 2020)**. Consistent with our study, decreased PAK3 impairs cognitive function via regulation of dendritic spine in neurodegenerative diseases. In addition to dendritic spine morphology regulation, PAK3 involves in synaptic transmission and trafficking. For example, PAK3 controls the surface trafficking of the major excitatory receptor GluA1 AMPA receptor subunit in neurons **(Hussain et al., 2015)**. PAK3 can also control synaptic GABA(A)R surface stability via G protein-coupled receptor kinase interacting ArfGAP 1/βPIX signaling **(Smith et al., 2014)**. These results suggest that PAK3 plays a key role in regulating synaptic function as well as the dendritic spine morphology.

Previous studies have shown that the activation of PAK3 is primarily regulated by phosphorylation **(Liu et al., 2021; Wu & Jiang, 2022)**, and phosphorylated PAK3 is correlated to activity of AMPA receptor **(Hussain et al., 2015)**, neuroplasticity **(Kim et al., 2017)**, spine morphogenesis **(Zhang et al., 2005)**, and neurite outgrowth **(Shin et al., 2006)**. On the contrary, our study demonstrated that a decrease in PAK3 expression alone can significantly alter spine morphogenesis. This finding suggests that PAK3 expression levels may be a critical determinant of dendritic spine development and synaptic plasticity, even regardless of the phosphorylation status. Furthermore, our study identifies a potential mechanism underlying the cognitive impairment associated with cranial irradiation, which downregulates PAK3 expression. These results highlight the importance of PAK3 expression in cognitive function and provide a potential therapeutic target for reducing the adverse effects of cranial irradiation on brain function.

The dendritic spine is one of the major factors influencing cognitive function. In our study, we observed changes in dendritic spines due to radiation exposure, followed by subsequent cognitive impairment. Additionally, we established that regulating PAK3, which affects dendritic spine maturation, can modulate radiation-induced cognitive dysfunction. However, considering that radiation can impact the entire nervous system and that neural circuit function, neuroinflammation, and astrogliosis can also influence cognitive function **(Makale et al., 2017)**, future studies is needed to investigate the mechanisms of factors beyond dendritic spine changes caused by radiation.

Intranasal administration can bypass blood-brain barrier and rapidly deliver drugs from the nasal mucosa to the brain with a non-invasive way and reducing systemic side effects **(Corrigan et al., 2015; Tucker et al., 2018)**. Recent studies reported that intranasal insulin administration for the treatment of CNS disorders including mild cognitive impairment and AD (ClinicalTrials.gov Identifier: NCT01767909, **(Craft et al., 2020)**). Additionally, previous studies reported that intranasal administration of microRNA is effective to ischemic brain injury **(Lawson et al., 2022)**, depression **(Guan et al., 2021)**, and AD **(Mai et al., 2019)**. In this study, we demonstrated the potential utility of intranasal administration of antagomiR-206-3p for restoring PAK3 levels and cognitive function in irradiated mouse models with marginal side effects. Other advantages of intranasal administration include avoiding the gastrointestinal tract or hepatic metabolism **(Mistry et al., 2009)**, and enhancing drug bioavailability and stability **(Quintana et al., 2016)**. A previous study reported that drug absorption through nasal cavity is more efficient than through gastrointestinal tract **(Furubayashi et al., 2007)**. In addition, intranasal administration of biomacromolecular agents, including nucleic acid-based drugs, remarkably reduced the degradation rate and increased the delivery efficiency compared to the oral route **(Fortuna et al., 2014)**. Collectively, intranasal administration provides a promising avenue for targeting and delivery to the brain with little adverse effects.

Overall, our study highlights the detrimental effects of cranial irradiation on cognitive function and provides new insights into the role of PAK3 in dendritic spine development and cognitive function. We provide the underlying mechanisms of PAK3-associated cognitive impairment after radiotherapy using both differentiated human neurons and mouse models. Considering the high delivery efficiency and the anti-cognitive impairment effects of antagomiR-206-3p from our preclinical results, intranasal administration of antagomiR-206-3p may offer a promising approach for reducing the adverse effects of PCI.

## Materials and methods

### Mice

Male mice, aged 7 weeks, were used in this study. All experiments were performed in accordance with the provisions of the NIH Guide for the Care and Use of Laboratory Animals. The mice were housed individually or in groups of up to five in sterile cages, and were maintained in animal care facilities in a temperature regulated room (23 ± 1℃) with a 12 h light–dark cycle. All animals were fed water and standard mouse chow *ad libitum*.

### Irradiation

All mice, including those in the sham (0 Gy) group, were anesthetized with an intraperitoneal (i.p.) injection of zoletil (5 mg/10 g) daily for 5 days. After sufficient anesthesia was induced, the mice were attached to a fixed plate, and irradiation was applied at locations ranging from 0.3 cm behind the eyes to the posterior part of both ears to ensure that the radiation surface did not involve the eyes, mouth, and spinal cord (to prevent the normal exercise ability of the mouse from being affected by irradiation). The brain of mice were irradiated with 2 Gy daily for five days at a dose rate of 600 MU/min using a TrueBeam STx.

### Behavior: Open field test

Depressive-like behavior of mice were assessed in rectangular chambers (Width x Length x Depth = 50 cm × 50 cm x 38 cm). Mice were habituated for 3 min in the chamber (without recording) then placed for another 5 min in center of the chamber (with recording). Localization was recorded and analyzed by the time in center zone using Noldus EthoVision XT software (Noldus Information Technology, Leesburg, VA).

### Behavior: Y-maze

Spontaneous alteration was measured using a Y-maze apparatus (three grey opaque plastic, symmetrical arm, length: 40 cm, arm bottom width: 3 cm, arm upper width: 13 cm, height of wall: 15 cm, angled at 120°, Busan, South Korea). Each mouse was placed in the central area. The number of entries into the arms and alterations were recorded for 10 min with the EthoVisionXT video-imaging system. Max alternation was defined as the number of possible alternations counted by the total number of arms entered minus 2. Alternation was defined as consecutive entries into 3 different arms and counted only if a mouse entered into the 3 arms of maze (without revisiting the first arm at the third visit) **(Hölter et al., 2015)**.

Spatial memory was tested by placing the test mouse into the Y-maze with one arm of the maze closed off during training (Training session). This arm is designated as the novel arm. After 4 hours, during which the mouse is removed from the maze, the mouse is placed back into the maze with the blockage removed (Testing session). The number of entries into the novel arm was recorded for 10 min with the EthoVisionXT video-imaging system **(Kraeuter et al., 2019)**.

### Behavior: Novel object recognition (NOR)

NOR assay was used to explore cognitive performance in the long-term memory in a 50 × 50 cm opaque square arena. On Day 1, the mice were habituated to an open field arena for 5 min. On Day 2, the mice were exposed to 2 identical objects for a 5-min period. On Day 3, one of the objects was replaced with a new object, and the mice were let to explore the arena for 5 min. Each trial was recorded and the videos were automatically analyzed by EthoVision XT software (Noldus). The time spent with the familiar and novel objects was measured **(Rungratanawanich et al., 2019)**.

### Intranasal administration

According to the manufacturer’s instructions and previous study **(Zhou et al., 2021)**, 40 nmol of antagomiR-206-3p (sequence: 5’-CCACACACUUCCUUACAUUCCA -3’) or antagomiR-NC (the antagomiR negative control, its antisense chain sequence: 5’-UCUACUCUUUCUAGGAGGUUGUGA-3’) was dissolved in 1 mL of RNase-free water. A total of 24 μL of the solution (1 nmol per one mouse) was instilled with a pipette, alternately into the left and right nostrils (1 μL/time), with an interval of 3–5 min. The respiration of mice was not blocked, and other disorders were not observed; all mice in different groups survived. The same dosage and mode of administration were used for intranasal Cy5-antagomiR-206-3p administration (n = 3) to trace the distribution of antagomiR-206-3p after administration. We observed the overall and its sagittal section of the brain using ChemiDoc at 24 h after intranasal administration of Cy5-antagomiR-206-3p.

### Dissection of prefrontal cortex and hippocampus

The dissection of mouse brain regions was performed following a previous study **(Spijker, 2011)**. First, to obtain the hippocampal region, we gently held the brain and opened the forceps, slowly separating the cortical halves. Once an opening had been created along the midline for approximately 60%, we directed the forceps (in the closed position) counterclockwise by 30–40° to expose the left cortex from the hippocampus, repeatedly opening the forceps as necessary. We then repeated the same procedure for the right cortex by pointing the forceps in a 30–40° clockwise direction until the upper part of the hippocampus became visible. At the most caudal part of the hippocampus/cortex boundary, we moved the small forceps through the cortex and used them to separate the hippocampus from the fornix. After removing the hippocampus, we used the large forceps to fold the cortex back into its original position. Subsequently, we placed the brain with the dorsal side and cut coronal sections to reveal the prefrontal cortex and striatum at different levels. Using a sharp razor blade, we made the first cut to remove the olfactory bulb and cut the section containing the prefrontal cortex.

### Primary neuronal cell culture

As previous study reported **(Kim et al., 2022)**, pregnant C57BL/6 mice were obtained from a specific pathogen-free colony at Oriental Inc. (Seoul, Korea). Embryos were dissected from embryonic day 15.5 mice and prepared for culturing. The prefrontal cerebral cortices were dissected under sterile conditions, and the meninges were pulled off. As previous study, the brain was cut into 200-400 mm sections, starting from the olfactory bulb **(Hilgenberg & Smith, 2007)**. Once the blade enters the cortex, begin cutting 600µm coronal sections. The appropriate slices obtained after sectioning were used for the primary culture. The prefrontal cerebral cortices were then digested with 2.5% Trypsin-EDTA (Gibco BRL) in DMEM media for 15 min at 37 ◦C. After digestion, cells were dispersed through a pipette and filtered through a 70 μm nylon net filter. The cells were then centrifuged 5min at 500g; washed with Neurobasal medium containing B27 serumfree supplement (Thermo Scientific) and penicillin/streptomycin (Thermo Scientific). The cultured cells were incubated at 37 ◦C in a humidified atmosphere of 5% CO2. In the analysis of molecular alterations, cultured neurons were sampled 4 days after irradiation.

### Image analysis

Primary cultured neuron and differentiated neuron was attached by a poly-d lysine–coated and laminin-coated confocal plate, respectively. Cells were fixed in 4% paraformaldehyde (5 min) and permeabilized with 0.1% Triton X-100 in PBS with 5% BSA (10 min). After blocking with 1% BSA in PBS (30 min), primary antibodies were incubated overnight at 4 ◦C and stained with secondary antibodies for 1 h. Coverslips were mounted with Fluoromount™ aqueous mounting medium. Samples were imaged with a confocal microscope (LSM 800 system). The microscopes were controlled by Zeiss black edition software. Secondary antibodies were labeled with Alexa Fluor 488, or 594.

### Dendritic spine distribution

The images of entire neurons were acquired at 20X magnification. For spine counts, the images were analyzed using a semiautomated analysis software (NeuronJ) publicly available at https://imagescience.org/meijering/software/neuronj/manual/ **(Zhang et al., 2022)**. Whole dendrite was subdivided into segments of 1 μm and number of spines across whole thickness were traced for length and breadth of each spine. Length and breadth ratio was used to determine the spine subtype as described earlier **(Bączyńska et al., 2021)**. After the analysis for each class of spine, standard deviation and p-values are calculated using two-way Anova with Sidak’s multiple comparison method.

### Real-time qRT-PCR

The expression levels of mRNAs, miRNAs, and stemness and neuron-related genes were analyzed by real-time qRT-PCR, as previously described **(Shin et al., 2022)**. Aliquots of the master mix containing all of the reaction components with the primers were dispensed into a real-time PCR plate (Applied Biosystems, Foster City, CA, USA). All PCR reagents were from a SYBR Green core reagent kit (Applied Biosystems). The expressions of all genes were measured in triplicate in the reaction plate. The qRT-PCR was performed using the StepOne Real-Time PCR System (Applied Biosystems). It was performed by subjecting the samples at 95 ◦C for 15 s and at 60 ◦C for 1 min for 40 cycles followed by thermal denaturation. The expression of each gene relative to Gapdh mRNA was determined using the 2^−ΔΔCT^ method.

### F/G-actin ratio

F/G-actin ratio was assessed as previously described **(Festa et al., 2020)**. Briefly, cells were lysed in cold lysis buffer [10 mM K_2_PO_4_, 100 mM NaF, 50 mM KCl, 2 mM MgCl_2_, 1 mM EGTA, 0.2 mM DTT, 0.5% Triton-X 100, 1 mM sucrose (pH 7.0)] and centrifuged at 15,000 x g for 30 min. Separation of F-actin and G-actin was achieved in that F-actin is insoluble (pellet) in this buffer, whereas G-actin is soluble (supernatant). The G-actin supernatant was transferred to a fresh tube and the F-actin pellet was resuspended in lysis buffer plus an equal volume of a second buffer [1.5 mM guanidine hydrochloride, 1 mM sodium acetate, 1 mM CaCl_2_, 1 mM ATP, 20 mM tris-HCl (pH 7.5)] and then incubated on ice for one hour with gentle mixing every 15 min to convert F-actin into soluble G-actin. Samples were centrifuged at 15,000 x g for 30 min and the supernatant (containing F-actin which was converted to G-actin) was transferred to a fresh tube. F-actin and G-actin samples were loaded with equal volumes and analyzed via Western blot.

### Western blot analysis

The assessment of protein levels was performed following previous study **(Kang et al., 2023)**. In brief, Whole cell lysates (WCL) were prepared using radioimmunoprecipitation assay (RIPA) lysis buffer (50 mM Tris, pH 7.4, 150 mM NaCl, 1% Triton X-100, 25 mM NaF, 1 mM dithiothreitol (DTT), 20 mM EGTA, 1 mM Na_3_VO_4_, 0.3 mM phenylmethanesulfonyl fluoride (PMSF), and 5 U/mL aprotinin) and the protein concentrations in the lysates were determined using a BioRad protein assay kit (BioRad Laboratories, Hercules, CA). Protein samples were subjected to SDS-PAGE, transferred to a nitrocellulose membrane and then blocked with 5% bovine serum albumin in TBST for 1 h at RT. Next, membranes were probed with specific primary antibodies and peroxidase-conjugated secondary antibody (Santa Cruz Biotechnology). The samples were subsequently analyzed using an ECL detection system (Roche Applied Science, Indianapolis, IN). Detection and densitometric analysis were conducted using the iBright Western blot imaging systems (Thermofisher).

### Statistical analysis

All numeric data are presented as the means ± standard deviation (SD) from at least three independent experiments. Experimental results were analyzed by one-way ANOVA for ranked data followed by Tukey’s honestly significant difference test. The Prism 9 software (GraphPad Software, SanDiego, CA) was used to conduct all statistical analyses. A p value < 0.05 was considered to be statistically significant.

## Availability of data and materials

The data that support the findings of this study are available from the corresponding author upon reasonable request.

## Ethics approval and consent to participate

The animal protocol used in this study was approved by the Pusan National University Institutional Animal Care and Use Committee (PNU-IACUC) for ethical procedures and scientific care (Approval Number PNU-2021-0055).

## Consent for publication

All co-authors have read and approved of its submission to this journal.

