## Supplementary Information for "PAK3 downregulation induces cognitive impairment following cranial irradiation"

### Supplemental Tables

**S1 Table.** Primers for determining levels of gene expression

### Supplemental Figures

**S1 Fig.** Cranial radiation affects depressive-like behavior (related to Figure 1)

**S2 Fig.** PAK3 is downregulated by IR in the PFC and hippocampus regions (related to Figure 2)

**S3 Fig.** OP18 and TBCB signaling are not affected by IR (related to Figure 2)

**S4 Fig.** Neural progenitor cell is differentiated to human neuron (related to Figure 4)

**S5 Fig.** miR-206-3p affects only PSD-95 but not other neuronal and non-neuronal markers (related to Figure 5)

**S6 Fig.** AntagomiR-206-3p attenuates depressive-like behavior (related to Figure 6)

**S7 Fig.** Summary of alterations to PAK3 signaling in dendritic spine upon IR exposure

**S1 Table. Primers for determining levels of gene expression**

| Gene name | Forward primer | Reverse primer |
|---|---|---|
| *PAK3* (mouse) | 5′- TTGGATAACGAAGAAAAACCCCC -3′ | 5′- GGGCACATCTGTGAGCCATAG -3′ |
| *PAK3* (human) | 5’- CAGCAGAAAGGGAACTATAGACACAT -3’ | 5’- GGCAAGCATAAAGCCTATGGAA -3’ |
| *GAPDH* (mouse) | 5’- GCAGTGTCATTAGCTGAT –3’ | 5′- CAGGACTTGCAATGTTGCTTGAG -3′ |
| *GAPDH* (human) | 5’- GGACGGGACGCGGTGCAG -3’ | 5′- CTTATAAGGCGCGGAACCGAAA -3′ |
| *SOX2* | 5’- TCCCCAGATACAATGGAC -3’ | 5′- TCCATGCTGTTTCTTACTCTCC -3′ |
| *NESTIN* | 5’- AGCCCTGACCACTCCAGTTTAG -3’ | 5′- CCCTCTATGGCTGTTTCTTTCTCT -3′ |
| *PAX6* | 5’- GCAACAACAGCAGCACAAAAA -3’ | 5’ - GGTTGTCACAGCTTCTGTCAAGA -3’ |
| *NeuN* | 5'- GGCGACCTACAGCATTGGA -3' | 5'- CATGGTCCGAGAAGGAAACG -3' |
| *PSD-95* | 5'- CGACAGCATCCTGTTTGTAAATG -3' | 5'- TCCACCGCCGCTGAGT -3' |
| *IBA-1* | 5'- CTGAAGGCCCAGCAGGAA -3' | 5'- TTTGGGATCGTCTAGGAATTGC -3' |
| *GFAP* | 5'- CGGCTGCTTTTCCCTAAGC -3' | 5'- GGGTACATTTTGTGTGTGAGTAAGAAG -3' |
| *miR-206-3p* | 5’- GCAGTGGAATGTAAGGAAGT -3’ | Universal 5’ - AGTGCAGGGTCCGAGGTATTC -3’ |
| *miR-1a-3p* | 5’- CGCAGTGGAATGTAAAGAAG -3’ | Universal |
| *miR-26a-5p* | 5’- GCAGTTCAAGTAATCCAGGATAG -3’ | Universal |

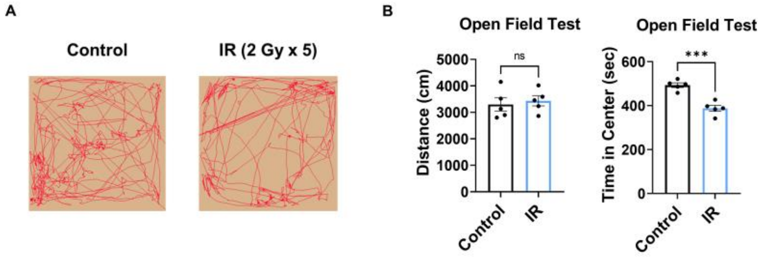


**S1 Fig. Cranial radiation affects depressive-like behavior (related to Figure 1)**

**(A)** The representative heatmaps of open field test after IR. (**B)** The result of open field test after IR. Statistical analysis was performed with Student's t test. ns, non-significant; ***$p < 0.001$.

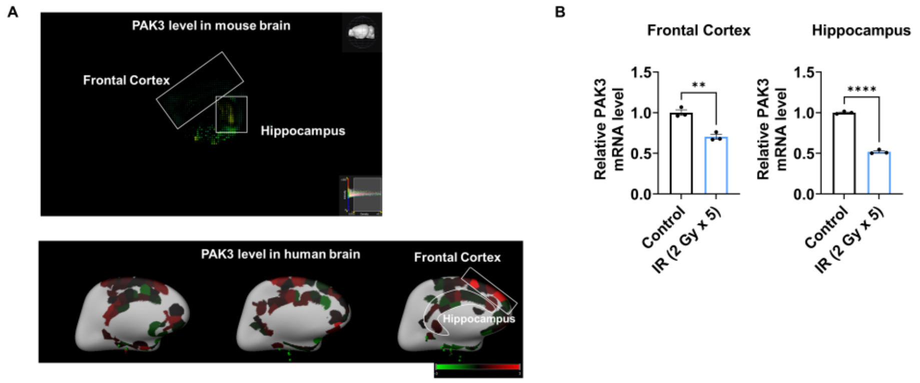


**S2 Fig. PAK3 is related to synapse organization and dendritic spine development in frontal cortex and hippocampus (related to Figure 2)**

**(A)** The localization of PAK3 in mouse and three human brain from Allen brain map (https://portal.brain-map.org/). **(B)** The changes of PAK3 mRNA level after IR in frontal cortex and hippocampus. Statistical analysis was performed with Student's t test. ***$p < 0.01$*; ****$p$ < 0.0001.

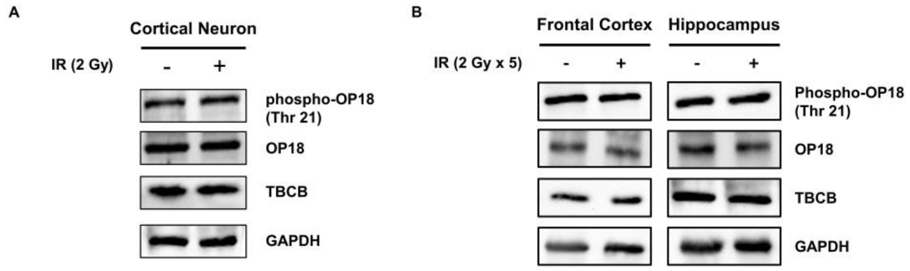


**S3 Fig. OP18 and TBCB signaling are not affected by IR (related to Figure 2)**

**(A)** The protein levels of phosphorylated OP18, OP18, TBCB after IR in cortical neuron. (**B)** The protein levels of phosphorylated OP18, OP18, TBCB after IR in frontal cortex and hippocampus.

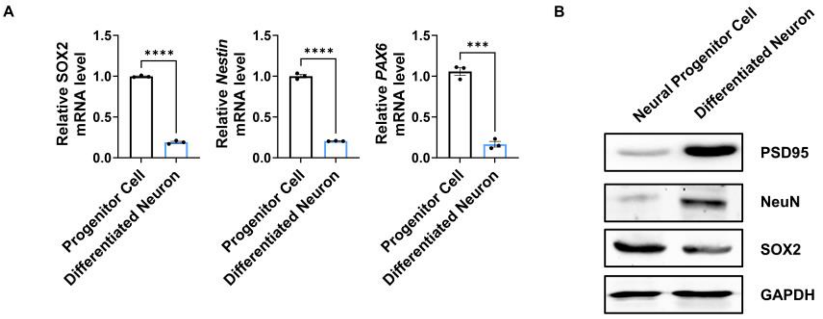


**S4 Fig. Neural progenitor cell is differentiated to human neuron (related to Figure 4)**

**(A)** The mRNA levels of SOX2, Nestin, and PAX6 in neural progenitor cell and differentiated neuron. (**B)** The protein levels of PSD95, NeuN, SOX2, and GAPDH. Statistical analysis was performed with Student's t test. ***$p$ < 0.001; ****$p$ < 0.0001.

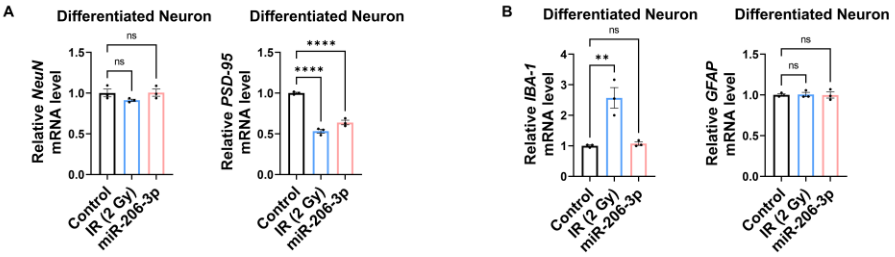


**S5 Fig. miR-206-3p affects only PSD-95 but not other neuronal and non-neuronal markers (related to Figure 5)**

**(A)** The mRNA levels of NeuN and PSD-95 after IR 2 Gy or miR-206-3p in differentiated neuron. **(B)** The mRNA levels of IBA-1 and GFAP after IR 2 Gy or miR-206-3p in differentiated neuron. Statistical analysis was performed with one-way ANOVA plus a Tukey's multiple comparisons test. ns, non-significant; $^{**}p < 0.01$; $^{****}p < 0.0001$.

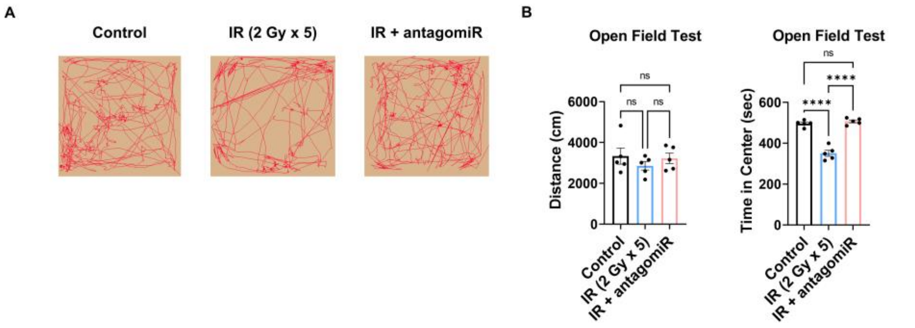


**S6 Fig. AntagomiR-206-3p attenuates depressive-like behavior (related to Figure 6)**

**(A)** The representative heatmaps of open field test after IR or IR with antagomiR-206-3p. (**B)** The result of open field test after IR or IR with antagomiR-206-3p. Statistical analysis was performed with one-way ANOVA plus a Tukey's multiple comparisons test. ns, non-significant; ****$p$ < 0.0001.

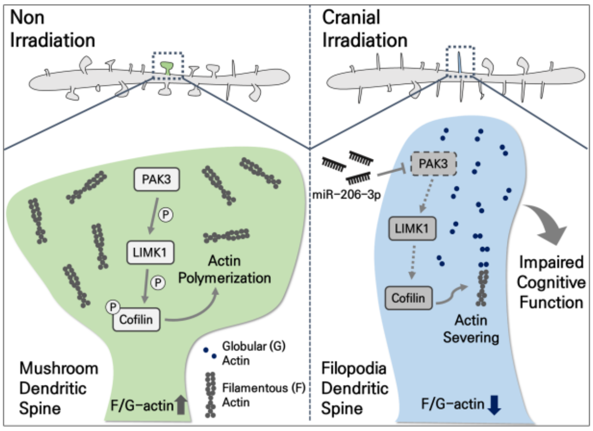


**S7 Fig. Summary of alterations to PAK3 signaling in dendritic spine upon IR exposure.**

Cranial irradiation impairs the maturation of dendritic spines by decreasing PAK3 signaling. Briefly, cranial irradiation increases miR-206-3p, which targets PAK3 and decreases its levels. Decreased PAK3 reduces the phosphorylation of LIMK and cofilin, resulting in decreased actin polymerization. This leads to a reduction in the ratio of F-actin to G-actin, which in turn impairs the maturation of dendritic spines. As a result, cognitive function is impaired via dendritic spines.
